# Targeting CCRL2 enhances therapeutic outcomes in a tuberculosis mouse model

**DOI:** 10.1101/2024.09.23.614576

**Authors:** Tianyin Wang, Darla Quijada, Taha Ahmenda, Jennie Ruelas Castillo, Nour Sabiha Naji, J David Peske, Petros C. Karakousis, Suman Paul, Theodoros Karantanos, Styliani Karanika

## Abstract

Tuberculosis (TB) remains among the leading infectious causes of death. Due to the limited number of antimicrobials in the TB drug discovery pipeline, interest has developed in host-directed approaches to improve TB treatment outcomes. C-C motif chemokine-like receptor 2 (CCRL2) is a unique seven-transmembrane domain receptor that is upregulated by inflammatory signals and mediates leucocyte migration. However, little is known about its role in the setting of TB infection. Here, we show that *Mycobacterium tuberculosis* (Mtb) infection increases CCRL2 protein expression in macrophages and in mouse lungs. To target selectively CCRL2-expressing cells *in vivo,* we developed a novel mouse anti-CCRL2 antibody-drug conjugate (ADC) linked with the cytotoxic drug SG3249. We tested its adjunctive therapeutic efficacy against TB when combined with the first-line regimen for drug-susceptible TB (isoniazid, rifampin, pyrazinamide, ethambutol; RHZE). The anti-CCRL2 ADC treatment potentiated RHZE efficacy in Mtb-infected mice and decreased gross lung inflammation. CCRL2 expression in lung dendritic cells and alveolar macrophages was lower in mice receiving anti-CCRL2 ADC treatment + RHZE compared to those receiving RHZE alone or the control group, although the total innate cell populations did not differ across treatment groups. Interestingly, neutrophils were completely absent in the anti-CCRL2 ADC treatment + RHZE group, unlike in the other treatment groups. IFN-γ+ and IL17-Α+ T-cell responses, which are associated with optimal TB control, were also elevated in the anti-CCRL2 ADC treatment + RHZE group. Collectively, our findings suggest that CCRL2-targeting approaches may improve TB treatment outcomes, possibly through selective killing of Mtb-infected innate immune cells.

## Introduction

Tuberculosis (TB) remains one of the leading infectious causes of death worldwide (WHO, 2023). The currently available six-month regimen consisting of isoniazid, rifampin, pyrazinamide, and ethambutol (collectively, RHZE) is highly effective against drug-susceptible TB. However, its length and complexity leads to treatment interruptions that adversely affect cure rates and increase the incidence of drug resistance (Fauci, 2008;Chuang et al., 2020). Thus, novel approaches that can shorten the duration of TB drug treatment are urgently needed to improve clinical outcomes in patients.

Due to the limited number of antimicrobials in the TB drug discovery pipeline, host-directed strategies have generated considerable interest (Kaufmann et al., 2018). Host immunity plays a critical role in the outcomes of TB disease (Young et al., 2020). *Mycobacterium tuberculosis* (Mtb) has developed several evasion strategies, prompting the host to elicit an immune response that favors its persistence (Young et al., 2020). Host-directed strategies targeted at “re-educating” the immune system are realistic alternative approaches to tailor the host response against TB (Young et al., 2020).

Neutrophils, macrophages, and dendritic cells (DCs) are among the major innate immune cell types involved in Mtb infection (Fulton et al., 2002;Eruslanov et al., 2005;Liu et al., 2017a). Although they are intended to target the invading pathogen, they may facilitate immune surveillance escape (Lee and Downey, 2001;Eruslanov et al., 2005;Liu et al., 2017b). Mtb-infected alveolar macrophages (AMs) have poor bactericidal capabilities and show immunoregulatory features suppressing lymphocyte activation (Guirado et al., 2013). Similarly, Mtb antigens alter differentiation, suppress maturation, and induce the expression of inhibitory molecules in DCs, suppressing their activity (Mihret, 2012;Magallanes-Puebla et al., 2018). Neutrophils display weak antimycobacterial activity and may promote mycobacterial survival through immune surveillance escape (Eruslanov et al., 2005). Neutrophil degranulation, which is meant to target Mtb, may also cause the destruction of neighboring cells and tissue dissolution (Fujie et al., 1999;Lee and Downey, 2001). Thus, further investigation of the molecular and cellular alterations mediating the phenotypic changes of these cells can improve our understanding of TB pathogenesis. Targeting molecules halting these alterations could be a promising approach to enhance the efficacy of currently available anti-TB therapies.

C-C motif chemokine-like receptor 2 (CCRL2) is a seven-transmembrane domain receptor that is upregulated by inflammatory signals (Schioppa et al., 2020). It does not promote chemotaxis but mediates leukocyte migration (Del Prete et al., 2017;Schioppa et al., 2020). CCRL2 has been recently found to promote malignant cell growth in myelodysplastic syndrome and secondary acute myeloid leukemia (Karantanos et al., 2022), while its deletion increased the sensitivity of these cells to azacytidine, a first-line chemotherapy for these malignancies (Karantanos et al., 2023). Data on the potential role of CCRL2 in TB are scarce. Petrilli et al. applied an immune-based gene expression profile and found that CCRL2 was one of the 7 genes that can predict the progression of latent TB infection to TB disease with high sensitivity and specificity (Petrilli et al., 2020). Consistently, silencing of CCRL2 abolished Mtb-induced macrophage M2 polarization, a phenotype associated with a suppressive microenvironment promoting Mtb intracellular growth (Zhang et al., 2020).

In this study, we show that Mtb infection increases the expression of CCRL2 *in vitro* and *in vivo,* and targeting CCRL2 with an antibody-drug conjugate (ADC) potentiates the bactericidal activity of RHZE in a mouse TB model, suggesting that this approach may be a promising adjunctive TB therapeutic tool.

## Materials and methods

### Bacteria

Wild-type and green fluorescent protein (GFP)-expressing Mtb H37Rv was grown in Middlebrook 7H9 broth (Difco, Sparks, MD) supplemented with 10% oleic acid-albumindextrose-catalase (OADC, Difco), 0.2% glycerol, and 0.05% Tween-80 at 37°C in a roller bottle (Karanika et al., 2022) .

### Cell line and *in vitro* reagents

Human THP1 cells were purchased from the American Type Culture Collection and cultured in RPMI-1640 with 10% heat-inactivated fetal bovine serum (hiFBS) with 2 mM l-glutamine, penicillin (100 U/ml), and streptomycin (100 μg/ml) at 37°C in 5% CO2. Phorbol 12-myristate 13-acetate (PMA) was purchased from Sigma-Aldrich (#P1585-5MG).

### Mtb infection *in vitro*

THP1 cells were maintained in RPMI 1640 supplemented with 10% hiFBS. Cells were differentiated with 50ng/ml PMA for 48 hours, washed, rested overight with media contaiing no PMA and infected in a growth medium containing 5% human serum. GFP+ bacteria were added to cells at a multiplicity of infection (MOI)= 1 or 10 for 1 or 2 days at 37°C, and extracellular bacteria were removed by sequential washes with PBS. Cells were then stained and analyzed by flow cytometry.

### Generation and analysis of antibody-drug conjugates

The anti-CCRL2 (BioLegend #114002 and #358304) and IgG2b (BioLegend # 400602 and 400608) mouse and human monoclonal antibodies were partially reduced using 5 molar excess Tris(2-carboxyethyl) phosphine hydrochloride (TCEP, Thermo Fisher Scientific, #77720) for 2 hours at 37°C with rotation, followed by TCEP removal by buffer exchange into PBS using Zeba Spin 7-kDa molecular weight cut-off (MWCO) columns. The SG3249 (Med Chem Express #HY-128952) solution was prepared by resuspending SG3249 in DMSO stock solution at 10 mM concentration. The SG3249 working solution was prepared by diluting the stock solution in sufficient DMSO to make the final conjugation reaction mixture 10% organic and 90% aqueous. This solution was added to the partially reduced antibodies at fivefold molar excess and incubated for 2 hours at room temperature with rotation. The excess unreacted drug linkers were removed by buffer exchange into PBS using Zeba Spin 7 kDa MWCO columns. The ADCs were analyzed for concentration using the Pierce BCA Protein Assay (Thermo Fisher Scientific, #23227), and drug–antibody conjugation and the presence of the residual free drug by high-performance liquid chromatography (HPLC) as previously described (Paul et al., 2021;Nichakawade et al., 2024).

### Mouse Mtb infections

All *in vivo* procedures were performed according to protocols approved by the Johns Hopkins University Institutional Animal Care and Use Committee. Four-to-six-week-old male and female C57BL6 mice were purchased from The Jackson Laboratory. The mice were housed in individually ventilated cages maintained on a 12:12h light/dark cycle with free access to food and water. They were monitored at least weekly, recording their weights and assessing their general appearance. They were infected between 6 and 8 weeks of age with ∼100 bacilli of wild-type Mtb H37Rv via the aerosol route using the Glas-Col Inhalation Exposure System (Terre Haute, IN), and this procedure did not require anesthesia. On the day after infection, at least 5 mice per experiment were sacrificed to determine the number of colony-forming units (CFUs_ implanted into the lungs.

### Mouse antibiotic treatments

When applicable, after 28 days of infection, at least 5 animals per experiment were sacrificed to determine the CFUs present in the lungs at the start of treatment. On the same day, the mice were randomized to receive either human-equivalent doses of H (10 mg/kg), R (10 mg/kg), Z (150 mg/kg), and E (100 mg/kg) dissolved in distilled water or distilled water only (control group) (Williams et al., 2009). The Z solution was gently heated in a 55°C water bath and vortexed to dissolve before treating mice. To minimize drug-drug interactions, R was administered at least 1 hour before Z. All drugs were administered once daily, by orogastric gavage, in a total volume of 0.2-0.3 ml/mouse/day.

### Mouse ADC treatments

Seven weeks after the Mtb aerosol infection, the CCRL2 and its isotope IgG control ADC were administered once, at a dose of 0.75 mg/kg, through a lateral tail vein injection using an insulin syringe with a 28-gauge needle. The needle was bent at an angle of 30–50°. The total volume did not exceed 100 μl. Animals were warmed under a heating lamp to promote vasodilatation for less than 30 seconds, and the needle was placed on the surface almost parallel to the vein and inserted carefully.

### Tissue collection and bacterial enumeration

Mice were sacrificed 7 weeks after drug treatment initiation (CCRL2 ADC experiment) or 16 weeks after Mtb infection (CCRL2 expression in Mtb-infected vs. uninfected mice). Right lower lungs were harvested and processed into single-cell suspensions after mechanical disruption and filtration through a 70-μm cell strainer and then additionally digested using collagenase Type I (ThermoFisher Scientific) and DNase (Sigma-Aldrich) for 20 min at 37°C (Karanika et al., 2022). Single-cell suspensions were resuspended in warm R10 (RPMI 1640 with 2 mM-glutamine, 100 U ml^−1^ penicillin, 100 μg ml^−1^ streptomycin, 20mM HEPES, 1% sodium pyruvate, 1% non-essential mino acids and 10% hiFBS; Atlantic Biologicals) for flow cytometry analysis, as detailed below. The right upper lung of each mouse was homogenized using glass homogenizers for bacterial burden assessment. Serial tenfold dilutions of lung homogenates in PBS were plated on 7H11 selective agar (BD) supplemented with 50μg/ml Cycloheximide, 50μg/ml Carbenicillin, 25μg/ml Polymixin B, and 20μg/ml Trimethoprim for CFU enumeration. Plates were incubated at 37°C and CFU were counted 4 weeks later by at least 2 investigators (Karanika et al., 2022). The left lung was fixed by immersion in 10% neutral buffered formalin for at least 48 hours, paraffin-embedded, and sectioned for immunohistochemical analysis.

### Immunohistochemical analyses

Following the preparation of paraffin lung sections, slides were prepared and stained using hematoxylin and eosin (H&E). The degree of inflammation in the lungs was quantified by measuring integrated density using the Image J software in various lung fields, at least 3 per mouse per group, by 2 investigators.

### Multiparameter flow cytometry

1x10^6^ THP-1 cells seeded for each replicate. Cells were collected with tryspin-EDTA and incubated for 5 minutes in incubator (5% CO2 at 37°C). Cells were resuspended in FACS buffer (PBS + 0.5% Bovine serum albumin (Sigma-Aldrich, St. Louis, MO) and stained with FcX Blocker for 5 minutes at room temperature. Then stained with Zombie NIR Fixable Viability Kit for 15 minutes at room temp. Then extracellular staining with the following anti-human antibodies: PE-conjugated anti-CCRL2 (Biolegend #358304), APC-conjugated anti-CD14 (Biolegend #325607), BV421-conjugated anti-CD11b (Biolegend #393113) for 30 minutes. Cells were fixed with 4% PFA for 30 minutes and then washed and resuspended with FACS buffer (PBS + 0.5% Bovine serum albumin (Sigma-Aldrich, St. Louis, MO). Single-cell suspensions from mouse lungs were prepared as above. For Intracellular Cytokine Staining (ICS), GolgiPlug cocktail (BD Pharmingen, San Diego, CA) was added for 4 hours and cells were collected using FACS buffer (PBS + 0.5% Bovine serum albumin (Sigma-Aldrich, St. Louis, MO). Intracellular proteins were stained for 20 minutes, and samples were washed, fixed and resuspended with FACS buffer. The following anti-mouse mAbs were used for ICS: PercPCy5.5 conjugated anti-CD3 (Biolegend Cat. No 100217), FITC-conjugated anti-CD4 (Biolegend Cat. No 100405), Alexa Fluor 700 conjugated anti-CD8 (Biolegend Cat. No. 155022), APC conjugated anti-IFN-γ, (Biolegend Cat. No. 505809), PE conjugated anti-IL-17α (Biolegend Cat. No 506903), BUV563-conjugated anti-CD19 (BD Biosciences Cat. No. 749028), BUV496-conjugated anti-NK-1.1 (BD Biosciences Cat. No. 741062), BV605-conjugated anti-Ly-6G (Biolegend Cat. No. 127639), BV421-conjugated anti-CD11c (Biolegend Cat. No. 117330), BV650-conjugated anti-CD64 (Biolegend Cat. No. 104732), BUV805-conjugated anti-CD11b (BD Biosciences Cat. No. 741934), BV510-conjugated anti-MHCII (Biolegend Cat. No. 107636), FITC-conjugated anti-CD103 (Biolegend Cat. No. 121420). BD Biosciences LSRII/Fortessa flow cytometers were used. Flow data were analyzed by FlowJo Software (FlowJo 10.9, LLC Ashland, OR).

### Statistics

All reported P values are from two-sided comparisons. Pairwise comparisons of group mean values for log_10_ CFU (microbiology data) and flow cytometry data were made using one-way analysis of variance followed by Tuckey’s or Fisher’s LSD multiple comparisons tests, or paired T-test, or Mann-Whitney test. All error bars represent the estimation of the standard error of the mean, and all midlines represent the group mean unless otherwise is specified. Prism 9.3 (GraphPad Software, Inc. San Diego, CA) was utilized for statistical analyses and figure generation. A significance level of α ≤ 0.05 was set for all experiments.

## RESULTS

### Mtb infection increases CCRL2 protein expression *in vitro* and *in vivo*

To evaluate the effect of Mtb infection on CCRL2 expression in macrophages *in vitro*, THP-1 cells were infected with a GFP-expressing Mtb strain following treatment with PMA to induce differentiation to macrophages (**Fig 1A**). Uninfected THP-1 cells were also included as controls. CD11b+ and CD14+ expression (measured as median fluorescent intensity) were used to confirm differentiation of THP-1 treated with PMA (**Supplementary** Fig. 1A). CCRL2 expression was significantly upregulated in alive Mtb-infected THP-1 cells compared to uninfected or GFP-cells on day 1 (P=0.023 between MOI 10 GFP+ vs. GFP-cells) (**Supplementary** Fig. 1B) and day 2 (P<0.001 between MOI 1 and 10 GFP+ vs. GFP-cells) (**Fig. 1B**).

**Figure 1.**
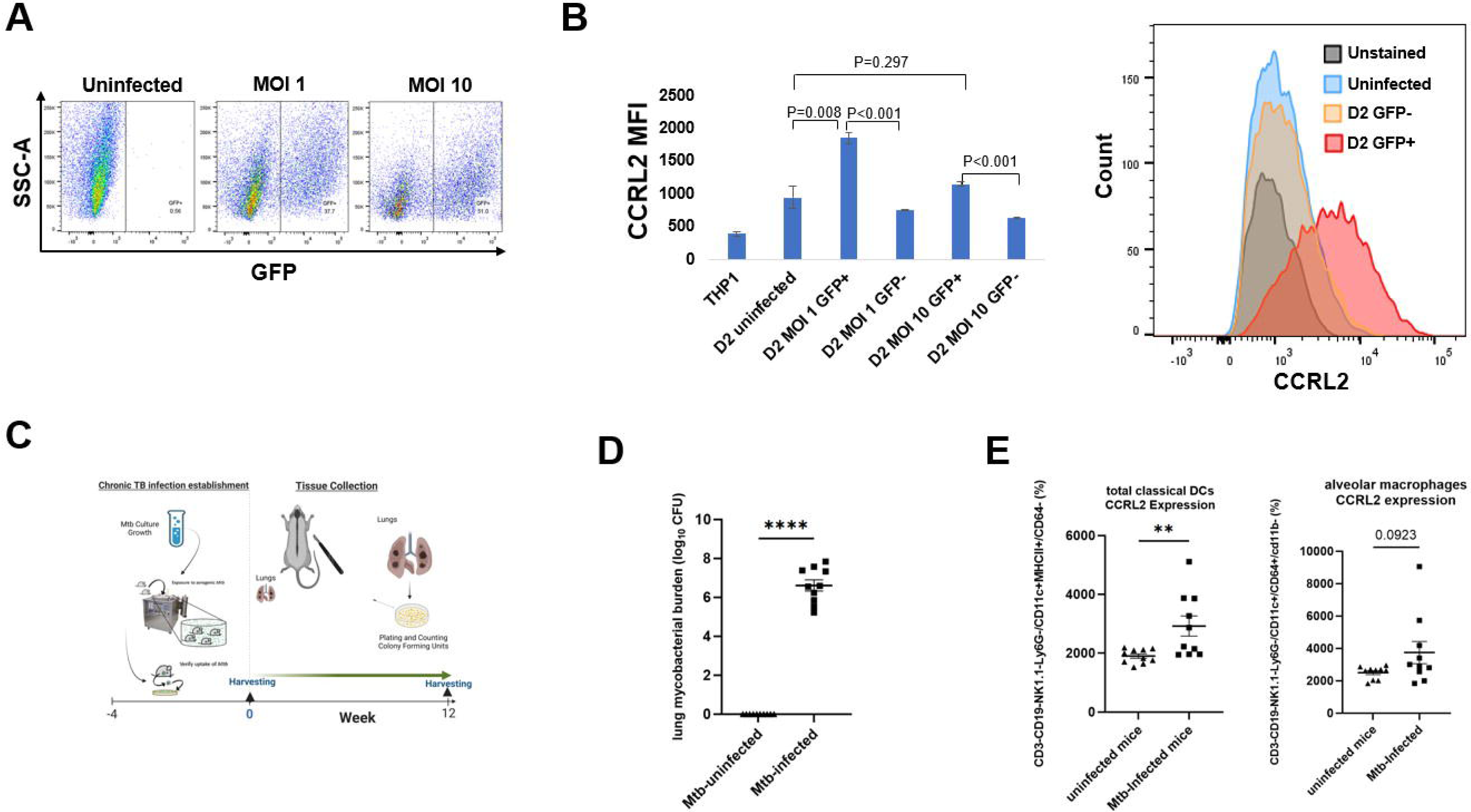
Mtb infection increases CCRL2 protein expression *in vitro* and *in vivo.* **(A),** **(B)** THP-1 cells were infected with the GFP+ Mtb strain at MOI of 1 and 10 following treatment with PMA to induce differentiation to macrophages and were incubated for 2 days. Using flow cytometry, CCRL2 expression (MFI) was measured across the different conditions; **(C)** Timeline of Mtb challenge study; **(D)** Mean lung mycobacterial burden between uninfected mice and Mtb infected mice at 12 weeks; and **(E)** CCRL2 expression of lung derived classical DCs and alveolar macrophages collected from Mtb-infected vs. uninfected mice. MOI, Multiplicity of infection; GFP, green fluorescent protein; D2, Day 2; MFI, Median fluorescent intensity; Mtb, *Mycobacterium tuberculosis*, CFU, colony-forming units; Classical DCs, classical dendritic cells; **P<0.01, ****P<0.0001

Next, to validate our *in vitro* findings, we sought to measure CCRL2 expression in the lungs of Mtb-infected and uninfected C57BL6 male and female mice. 16 weeks after infection, the lungs were collected (**Fig. 1C**) and uninfected mice of similar age were used as controls (**Fig. 1D**). CCRL2 expression was significantly higher in classical DCs in the lungs of infected mice compared to those of uninfected mice (P=0.009). A similar trend was also found in CCRL2 expression in alveolar macrophages (P=0.0923) (**Fig. 1E**).

### Treatment with an anti-CCRL2 ADC potentiates the activity of RHZE and decreases gross lung inflammation

Next, to investigate the effect of CCRL-2 deletion on TB treatment outcomes *in vivo*, we tested the adjunctive therapeutic efficacy of an anti-CCRL2 ADC when combined with the first-line regimen for drug-susceptible, RHZE. The mouse anti-CCRL2 ADC was generated by conjugating the commercially available anti-mouse CCRL2 antibody with the cytotoxic DNA minor groove inter-strand pyrrolobenzodiazepine dimer SG3249. An anti-mouse rat IgG2b was conjugated with SG3249 as an isotope control. The conjugation was confirmed by HPLC (**Fig. 2A and Supplementary Fig. 2A)**.

**Figure 2.**
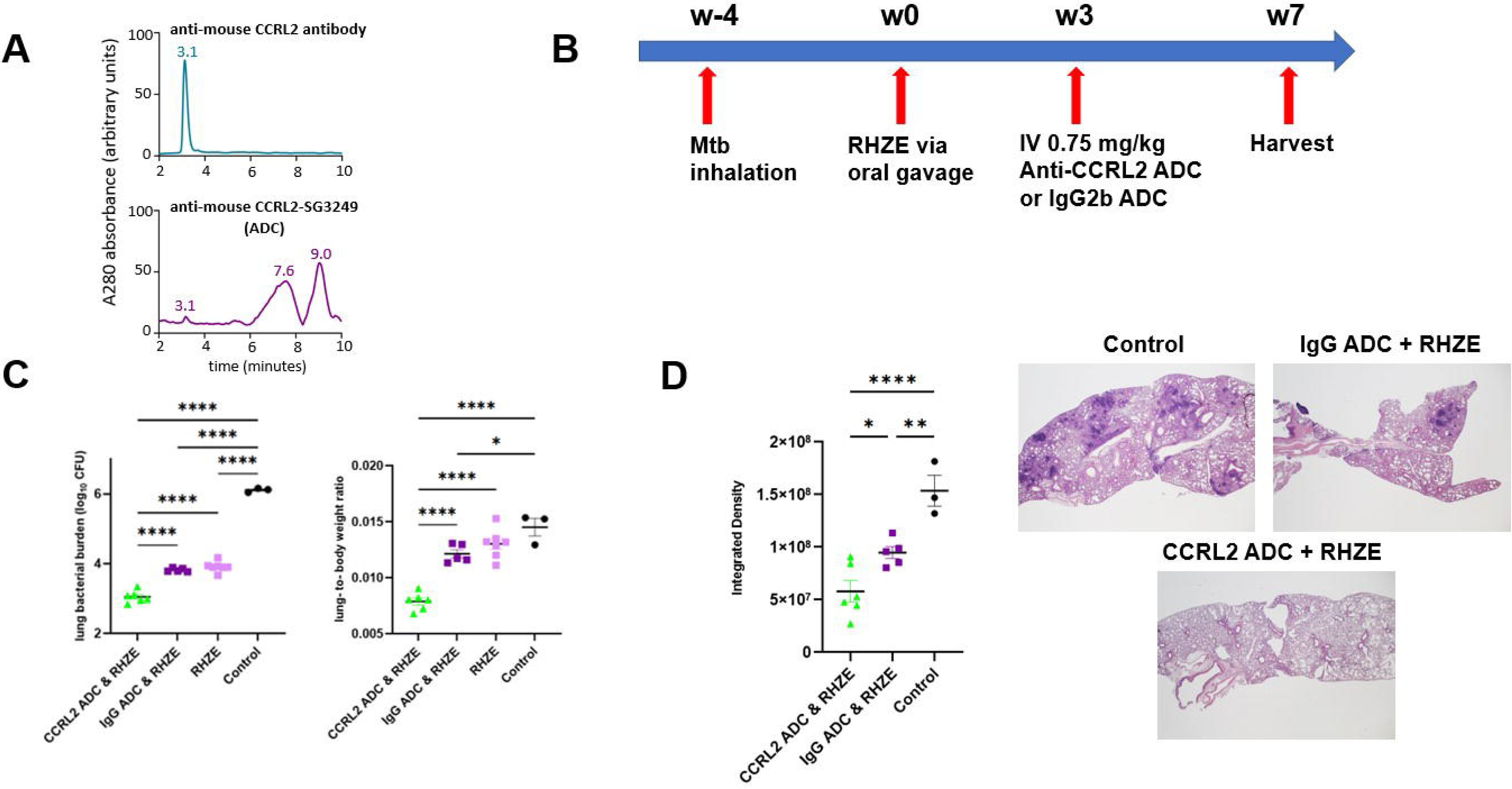
Treatment with an anti-CCRL2 ADC potentiates the activity of RHZE and decreases gross lung inflammation. **(A)** Hydrophobic interaction chromatography (HPLC) of anti-CCRL2 antibody, and anti-CCRL2-SG3249 ADC; **(B)** Timeline of the Mtb challenge study; **(C)** Scatterplot of mean lung mycobacterial burden in mice and lung-to body-weight ratio at 7 weeks after RHZE treatment initiation; **(D)** Scatterplot of integrated density, as measure of gross lung inflammation, across the different treatment groups and representative H & E slides per group; Mtb, *Mycobacterium tuberculosis*, RHZE, Rifampin-Isoniazid-Pyrazinamide-Ethambutol; ADC, Antibody-drug conjugate; CFU, colony-forming units;*P<0.01, **P<0.01, ****P<0.0001.

C57BL/6 mice were aerosol-infected with Mtb, and 4 weeks later, antibiotic treatment with RHZE was initiated (**Fig. 2B**). Three weeks later, the mice were intravenously treated with one dose (0.75 mg/kg) of the anti-CCRL2 ADC or the isotope control, anti-IgG ADC. Mice were sacrificed 4 weeks later to assess lung mycobacterial burden (**Fig. 2B**). The lung bacterial burden was found to be significantly lower in mice treated with anti-CCRL2 ADC + RHZE compared to those treated with the RHZE + isotope control (P<0.0001) or RHZE alone (P<0.0001) (**Fig. 2C**). The mouse lung-to-body weight ratio, an index reflective of gross lung inflammation, was significantly lower in mice treated with the anti-CCRL2 ADC than those treated with the anti-IgG ADC (P<0.0001) or RHZE alone (P<0.0001) (**Fig. 2C**). Of note, no toxicity was appreciated during the study in any of the treatment groups as measured by weight gain (**Supplementary Fig. 2B**), signs of morbidity or mortality. Similarly, inflammation was also evaluated histopathologically. Lung inflammation, as measured by surface area containing inflammatory lesions obliterating the lung parenchyma (integrated density), was significantly reduced in mice treated with adjunctive anti-CCRL2 ADC compared to the those treated with the anti-IgG ADC (P=0.015) **(Fig. 2D)**.

### Treatment with anti-CCRL2 ADC lowers CCRL2 expression in innate immune cells and induces TB-protective immune responses

To gain insight into the potential mechanisms underlying the adjunctive therapeutic efficacy of the anti-CCRL2 ADC, we measured the CCRL2 expression in innate immune cells derived from the lungs of mice receiving each of the different treatment groups using flow cytometry. We found that mice treated with anti-CCRL2 ADC + RHZE had nonsignigicantly lower CCRL2 expression (expressed in median fluorescent intensity) in lung classical DCs 1 (P=0.2366) and DCs 2 (P=0.0772) and alveolar macrophages (P=0.076) compared to those treated with the anti-IgG ADC + RHZE (**Fig. 3A**). Interestingly, the populations (%) of classical DCs 1 and 2 and alveolar macrophages isolated from the lungs of mice treated with anti-CCRL2 ADC + RHZE either were relatively increased or did not have significant variation compared to the IgG ADC + RHZE group (P=0.3716, 0.1390, 0.0895, respectively) **(Fig. 3B).** Notably, the neutrophils isolated from the lungs of mice treated with anti-CCRL2 ADC + RHZE were undetectable in contrast to the IgG ADC + RHZE or control group (P=0.0062 and <0.0001, respectively) (**Fig. 3B**). Taken together, these findings suggest that the adjunctive therapeutic efficacy of the CCRL2 ADC treatment may be associated with lower CCRL2 expression by DCs and alveolar macrophages, as well as a dramatic reduction of neutrophils.

**Figure 3.**
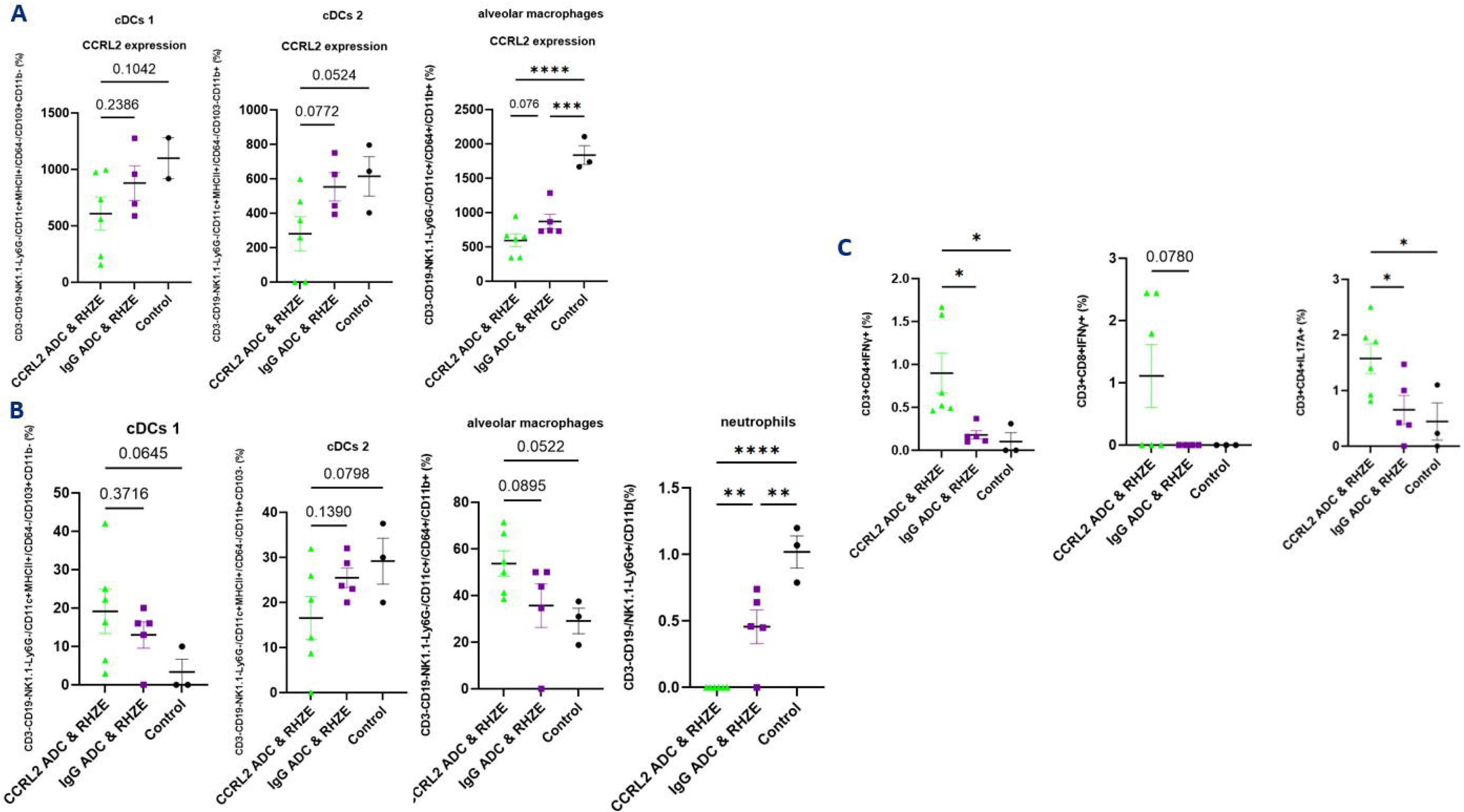
Treatment with an anti-CCRL2 ADC lowers CCRL2 expression in innate immune cells and induces TB-protective immune responses. **(A)** CCRL2 expression (MFI) in cDCs 1, cDCs 2, and alveolar macrophages derived from mouse lungs across the different treatment groups; **(B)** populations (%) of cDCs 1, cDCs 2, alveolar macrophages, and neutrophils derived from mouse lungs across; **(C)** IFNγ-CD4+ and CD8+ T cell producing cells and IL17Α-CD4+ T cell producing cells (%) across the different groups; Assessment was performed using flow cytometry; RHZE, Rifampin-Isoniazid-Pyrazinamide-Ethambutol; cDCs I, classical dendritic cells type I; cDCs II, classical dendritic cell type 2; ADC, antibody-drug conjugate. MFI, median fluorescent intensity.

Finally, we measured the T-cell immune responses associated with optimal TB control in the murine lungs (Khader et al., 2007;Cooper, 2009;Darrah et al., 2020;Karanika et al., 2024). The percentage of CD4+ and CD8+ T cells producing IFN-γ, and CD4+ T cells producing IL17-A were either relatively or significantly increased in the lungs of mice receiving anti-CCRL2 ADC + RHZE compared to those receiving anti-IgG ADC + RHZE (P=0.0122, P=0.078 and P=0.0305, respectively) (**Fig. 3C**).

## DISCUSSION

In this work, we show that CCRL2, a unique non-signaling seven-transmembrane domain receptor known to be upregulated in inflammatory signals (Del Prete et al., 2017;Schioppa et al., 2020;Zhang et al., 2020;Karantanos et al., 2022), may play an important role in TB pathogenesis. Specifically, we found that CCRL2 expression is increased in Mtb-infected differentiated THP-1 cells and also higher in lung-derived DCs and alveolar macrophages in Mtb-infected mice compared to uninfected mice. Targeting CCRL2 using an anti-CCRL2 ADC offered adjunctive therapeutic activity in Mtb-infected mice when combined with the first-line TB regimen. Anti-CCRL2 ADC treatment lowered CCRL2 expression by lung DCs and alveolar macrophages and abolished lung neutrophils. It also induced local TB-protective T-cell responses, suggesting that this novel host-directed therapy may be a promising therapeutic tool.

The role of CCRL2 in immunity is not clearly described. It is suggested that it responds to inflammatory signals and one of the proposed mechanisms of action involves the formation of heterodimers with chemokine receptors (Del Prete et al., 2017). Heterodimerization of chemokine receptors is a potentially crucial step for the proper function of immune cells (Mellado et al., 2001;Martínez-Muñoz et al., 2018). In particular, CCRL2/CXCR2 heterodimers control neutrophil recruitment in inflammatory arthritis (Del Prete et al., 2017), resulting in protection in CCRL2-deficient mice (Del Prete et al., 2017;Regan-Komito et al., 2017). Similarly, CCRL2 expression is upregulated in synovial neutrophils of patients with rheumatoid arthritis compared to healthy controls (Galligan et al., 2004). In an allergen-induced airway inflammation model, where DCs are known to play a crucial role (Lambrecht, 2005), CCRL2 deficiency impaired DC migration, leading to an adaptive immune response defect, which was abrogated with adoptive transfer of wild-type DCs (Otero et al., 2010). Silencing of CCRL2/CX3CR1 abolished Mtb-induced macrophage M2 polarization, a cell population associated with Mtb growth (Zhang et al., 2020). Taking together all of the available evidence, we hypothesized that TB, as an inflammatory disease, promotes CCRL2 expression. Indeed, we found that CCRL2 expression is higher in Mtb-infected differentiated THP-1 cells, and this was also confirmed *in vivo.* Adjunctive treatment with the novel targeted therapy anti-CCRL2-ADC further reduced the lung bacillary burden *in vivo*.

ADCs have been developed and successfully introduced in clinical practice as disease-selective treatments for various types of cancer, particularly hematologic malignancies (Ogitani et al., 2016;Zammarchi et al., 2018;Nichakawade et al., 2024;Paul et al., 2024). The delivery of the toxic drug selectively to cells expressing a surface marker allows disease-specific therapy while sparing other cells. CCRL2 is expressed at very low levels in healthy hematopoietic stem and progenitor cells (Karantanos et al., 2022) and at low levels in healthy non-inflamed tissues (Monnier et al., 2012;Del Prete et al., 2017). Of note, global deficiency of CCRL2 did not alter normal development and had no significant impact on blood counts or incidence of infection in mouse models (Del Prete et al., 2017;Xu et al., 2022). These findings render CCRL2 an attractive and safe target for selective therapies, including our novel anti-CCRL2 ADC, which had no associated toxicity when tested *in vivo*.

One possible mechanism of action of our anti-CCRL2 ADC is by selectively killing Mtb-infected innate immune cells, which contribute to immune surveillance escape. Indeed, we observed lower CCRL2 expression in lung DCs and alveolar macrophages of mice receiving the anti-CCRL2 ADC + RHZE, which was accompanied by the lowest lung mycobacterial burden compared to the anti-IgG ADC + RHZE or control groups, although the total population of DCs or alveolar macrophages was either unchanged or higher in the former group. Also, we observed that the neutrophil population was completely abolished in the anti-CCRL2 ADC + RHZE mouse group, suggesting that neutrophils are major contributors to the TB-associated inflammatory response and express CCRL2 targeted by the novel anti-CCRL2 ADC. This is not surprising since neutrophilia is independently associated with an increased risk of cavity formation, lung tissue damage, and mortality in patients undergoing TB therapy, suggesting that the neutrophil count in TB positively correlates with bacillary load and disease outcome (Lowe et al., 2013;de Melo et al., 2018).

Our study has some limitations. Although our findings allow us to make some initial associations between the adjunctive therapeutic role of the anti-CCRL2 ADC treatment and CCRL2 expression in the innate immune cells, further studies are needed to elucidate the mechanism of action in the TB setting. Also, additional studies are needed to test the therapeutic efficacy of this novel intervention in other animal models that more closely represent human TB pathology.

In conclusion, our data suggests that Mtb infection increases CCRL2 expression, and CCRL2-targeting approaches may improve TB treatment outcomes, possibly through selective killing of innate immune cells harboring Mtb. Ultimately, the potential utility of this novel strategy must be evaluated as an adjunctive therapeutic intervention in shortening the duration of curative treatment for active TB.

## Acknowledgements

Content of this publication has been available online in Biorxiv preprint server.. Illustrations were created with BioRender.com.

## Conflict of interest

The authors declare that the research was conducted in the absence of any commercial or financial relationships that could be construed as a potential conflict of interest. The Johns Hopkins University has filed patent applications related to technologies described in this paper on which SP, TK and SK are listed as inventors. Additional patent applications on the work described in this paper may be filed by Johns Hopkins University. The terms of all of these arrangements are being managed by Johns Hopkins University according to its conflict of interest policies. SP is a consultant to Merck, owns equity in Gilead and received payment from IQVIA and Curio Sciences.

## Data availability statement

The original contributions presented in the study are included in the article/Supplementary Material. Further inquiries can be directed to the corresponding authors.

## Ethics statement

The animal study was reviewed and approved by Johns Hopkins University Institutional Animal Care and Use Committee.

## Author contributions

SK and TK conceived and designed the study and wrote the manuscript. DG and TK performed *in vitro* studies. SK, TK, TW, DG, and JRC performed the *in vivo* experiments and their analysis. TA, SP, TN, NSN, SP and TK developed anti-CCRL2 and anti-IgG ADCs. JDP and TK performed immunohistochemistry analysis. SK, TK, TW, JDP performed data and statistical analysis. SK, TK, JDP, TW, DQ, PCK, JRC, DQ, TA, SP, TN, NSN, SP interpreted the data and edited the manuscript. All authors contributed to the article and approved the submitted version.

## Funding

This work was supported by NIH grants: K08AI174959 (SK), K08HL168777, K08CA270403 (SP) R01AI148710 (PCK), K24AI143447 (PCK), P30AI168436 (PCK). The content is solely the responsibility of the authors and does not necessarily represent the official views of the National Institutes of Health. SK was also supported by the Johns Hopkins CFAR developmental award (5P30AI094189-03) and Johns Hopkins Tuberculosis Research Advancement Center (TRAC) developmental award (P30AI168436). TK was also supported by the Leukemia Research Foundation. SP was also supported by the Leukemia Lymphoma Society Translation Research Program awards, the American Society of Hematology Scholar award and the Swim Across America Translational Cancer Research Award.

---

**Supplementary Figure 1. CCRL2 expression in differentiated Mtb-infected THP-1 cells; (A)** CD11b+ and CD14+ expression (measured as MFI through flow cytometry) were used to confirm differentiation of THP-1 into macrophages treated with PMA **(B)** THP-1 cells were infected with the GFP+ Mtb strain at MOI of 1 and 10 following treatment with PMA to induce differentiation to macrophages and were incubated for 1 day. Using flow cytometry, CCRL2 expression (MFI) was measured across the different conditions; MOI, Multiplicity of infection; GFP, green fluorescent protein; D1, Day 1; MFI, Median fluorescent intensity.

**Supplementary Figure 2. Hydrophobic interaction chromatography (HPLC) of isotype control antibody, and isotype control-SG3249 ADC. (A)** ADC, antibody-drug conjugate; **(B)** Mouse body weights (g) at 7 weeks after RHZE initiation.

