## Supplementary figures and images for "Targeting CCRL2 enhances therapeutic outcomes in a tuberculosis mouse model"

### Supplementary Figure 1

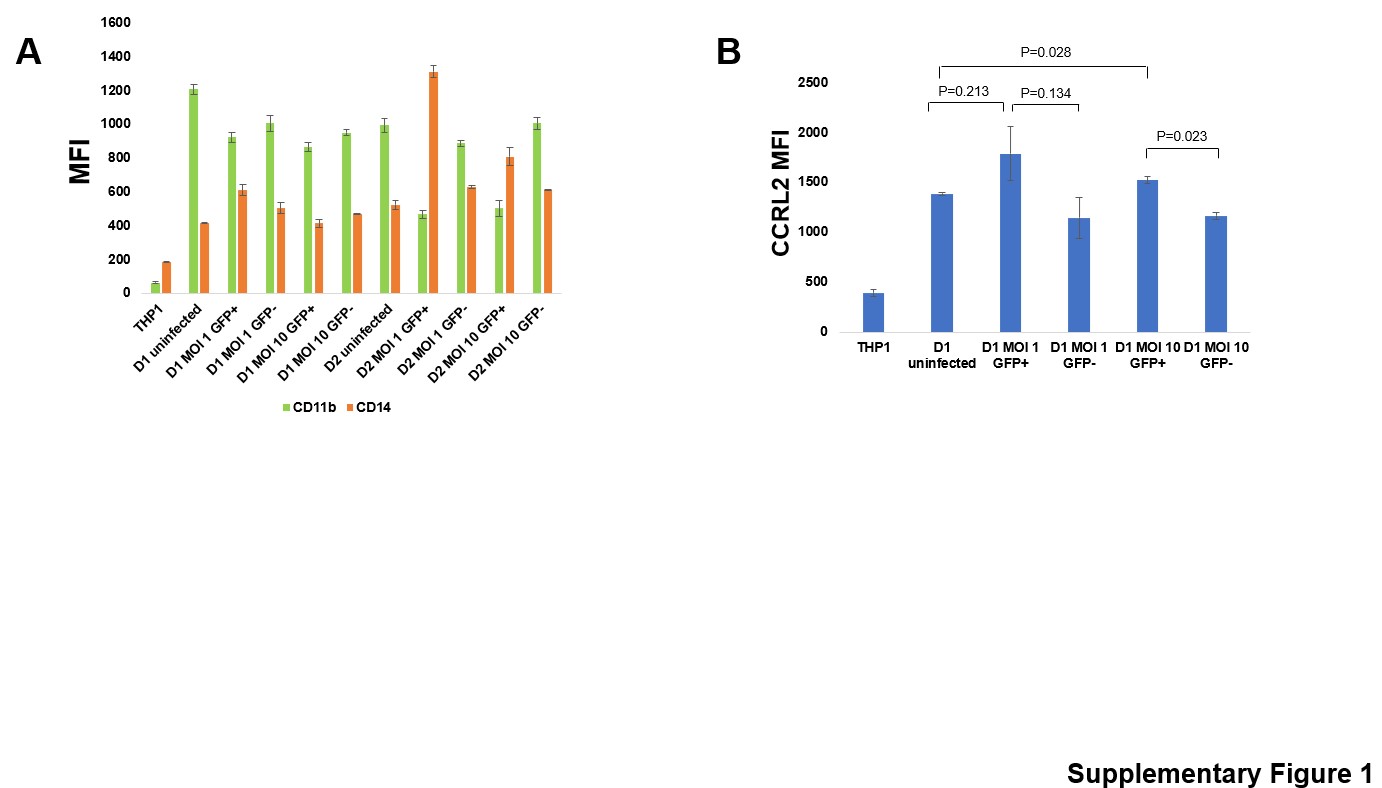

### Supplementary Figure 2

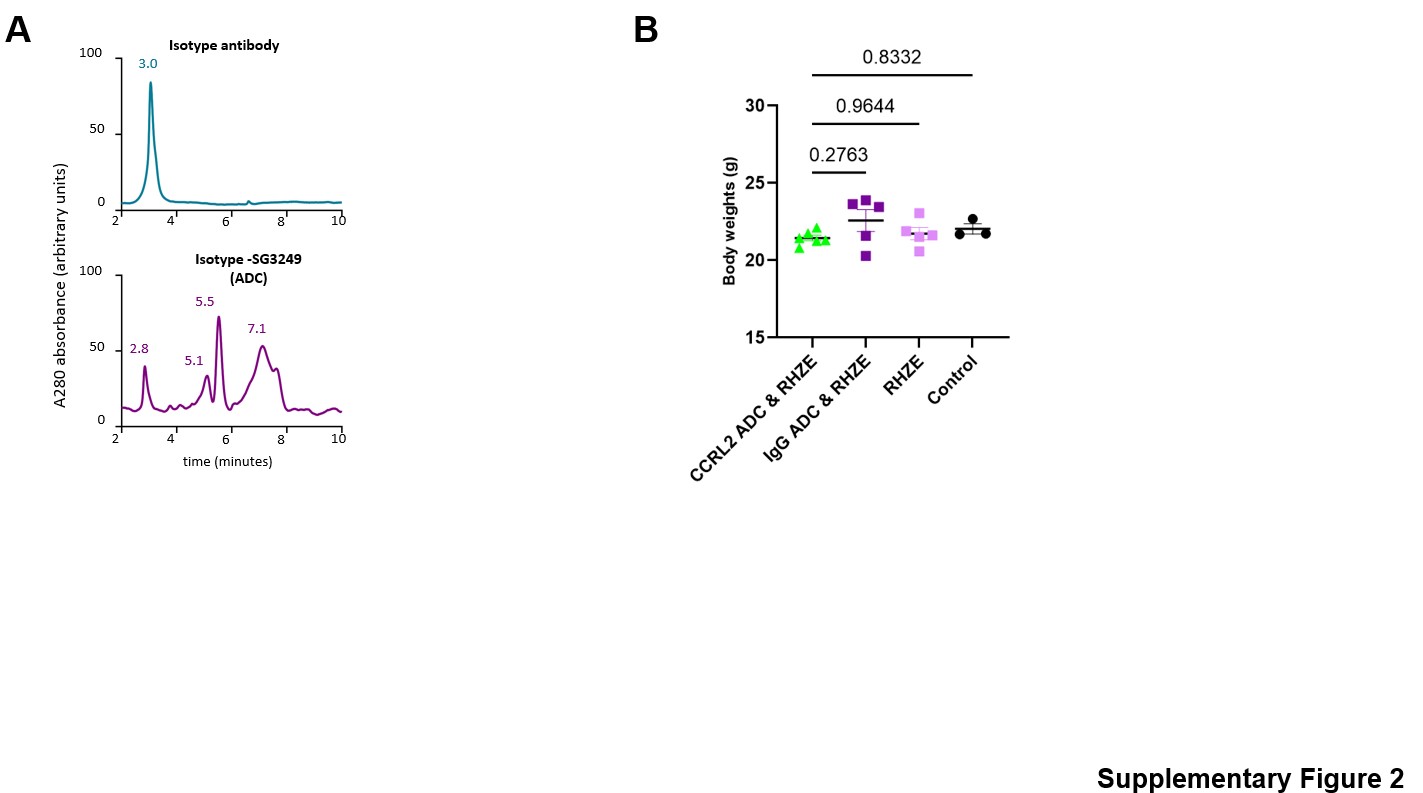
